# ^13^C tracer analysis reveals the landscape of metabolic checkpoints in human CD8^+^ T cell differentiation

**DOI:** 10.1101/2023.05.18.541159

**Authors:** Alexander Kirchmair, Niloofar Nemati, Giorgia Lamberti, Marcel P. Trefny, Anne Krogsdam, Anita Siller, Paul Hörtnagl, Petra Schumacher, Sieghart Sopper, Adolf M. Sandbichler, Alfred Zippelius, Bart Ghesquière, Zlatko Trajanoski

## Abstract

Naïve T cells remain in an actively maintained state of quiescence until activation by antigenic signals, upon which they start proliferation and generation of effector cells to initiate a functional immune response. Metabolic reprogramming is essential to meet the biosynthetic demands of the differentiation process, and failure to do so can promote the development of hypofunctional exhausted T cells. Here we used ^13^C metabolomics and transcriptomics to study the metabolic dynamics of CD8^+^ T cells in their complete course of differentiation from naïve over stem-like memory to effector cells. The quiescence of naïve T cells was evident in a profound suppression of glucose oxidation and a decreased expression of *ENO1*, downstream of which no glycolytic flux was detectable. Moreover, TCA cycle activity was low in naïve T cells and associated with a downregulation of SDH subunits. Upon stimulation and exit from quiescence, the initiation of cell growth and proliferation was accompanied by differential expression of T cell regulatory genes and metabolic reprogramming towards aerobic glycolysis with high rates of nutrient uptake, respiration and lactate production. High flux in anabolic pathways imposed a strain on NADH homeostasis, which coincided with engagement of the proline cycle for mitochondrial redox shuttling. With acquisition of effector functions, cells increasingly relied on glycolysis as opposed to oxidative phosphorylation, which paradoxically was not linked to changes in mitochondrial abundance. We further investigated the metabolic phenotype of exhausted T cells, finding that decreased effector function concurred with a reduction in mitochondrial metabolism, glycolysis and amino acid import, and an upregulation of suppressive and quiescence-associated genes, including *TXNIP* and *KLF2*. Thus, these results identify multiple features critical for the metabolic reprogramming that supports quiescence, proliferation and effector function of CD8^+^ T cells during differentiation. Further, an impairment of the same processes in exhaustion suggests that targeting these control points may be useful for both modulation of differentiation and prevention of exhaustion.

## Introduction

CD8^+^ T lymphocytes are cells of the adaptive immune system capable of acquiring cytotoxic functions. Following their development in the thymus, they persist in a quiescent state as naive T cells until activation by cognate antigen. This priming results in selective expansion of antigen-specific cells, which culminates in the mounting of an effective immune response. Due to their potentially disease-causing effects, cytotoxic cells require tight regulation and successful immune responses need to be rapidly terminated. Only a subset of antigen-specific cells survives long-term as memory cells for preparedness to antigen reencounter. Early studies revealed the presence of two major subpopulations of memory cells, central memory (TCM) cells that are primarily homed to lymph nodes and effector memory (TEM) cells with increased cytotoxic potential in the peripheral circulation (Sallusto et al. 1999). A rare population of stem cell memory cells (TSCMs) has later been identified in mice (Y. Zhang et al. 2005) and human studies (Gattinoni et al. 2011). These cells are capable of long-term persistence, generate memory responses to antigen reencounter and produce TCM and TEM offspring. Further lineage analyses in vivo confirmed that stem-like T cells also sustain the primary immune response against cancer (Prokhnevska et al. 2023), viral infections (Pais Ferreira et al. 2020), vaccines (Jung et al. 2022) and auto-immune targets (Gearty et al. 2022). Thus, while dedifferentiation of effector into memory cells may occur in some settings (Youngblood et al. 2017), the majority of studies supports a progressive “memory-first” differentiation model, where TSCM cells with long-term persistence generate more differentiated, short-lived progeny for effector responses. This differentiation process needs to be tightly regulated to control the magnitude, effectiveness and dynamics of an immune response, and various regulatory points evolved for this purpose (ElTanbouly and Noelle 2021). For example, the quiescence of naive T cells is enhanced by checkpoints such as VISTA (ElTanbouly et al. 2020), NRP1 restricts memory formation (Liu et al. 2020) and activation and effector functions are restrained by PD1 ligation (Freeman et al. 2000).

Conceivably, the biosynthetic and bioenergetic demands of distinct T cell subsets are reflected in their metabolic needs. T cell activation induces a switch to glycolysis (Gupta, Wang, and Chen 2020) to provide precursors for growth (Lunt and Vander Heiden 2011), whereas resting naive and memory cells mainly rely on fatty acid oxidation (FAO) and oxidative phosphorylation (OXPHOS) to meet their basal energy requirements (Corrado and Pearce 2022). Mitochondrial function has also been found to be impaired in and contribute to exhaustion (Schulz and Zehn 2022). These metabolic programs are sometimes connected to specific immunometabolites that exert signaling functions, such as phosphoenolpyruvate (Ho et al. 2015), asparagine (J. Wu et al. 2021), kynurenine (Opitz et al. 2011), succinate (Elia et al. 2022) or itaconate (Lin et al. 2021). In consequence, metabolic cues in the microenvironment have the potential to modulate T cell differentiation and may be co-opted not only by tumor cells, but also for cancer therapy. Of particular interest in anti-cancer immunity are checkpoints that impair cytotoxicity in exhausted T cells. Exhaustion presumably functions as a physiological adaptation to persistent antigen presence, but becomes a dysfunctional adaptation when tumor-associated microenvironmental signals result in reinforced immune suppression and subsequent failure to eradicate a tumor (Zarour 2016). Such metabolic dysregulation is characterized by decreased glycolysis, OXPHOS and ATP production, whereas oxidative stress and ROS levels are elevated. Overall, this results in a disability of exhausted T cells to accomplish their role as cytotoxic cells (Chan et al. 2022).

Moreover, it has become clear that the pathways regulating differentiation and exhaustion comprise more than classical signaling cascades and transcription factors, but also integrate metabolic systems. While many of the initial metabolic investigations focused on measuring metabolite concentrations, stable-isotope tracing is increasingly employed to yield insights into activities and fluxes of metabolic pathways in immune cells, e.g. (Ma et al. 2019), (Hermans et al. 2020), (J. Wu et al. 2021). However, metabolic flux dynamics have so far not been investigated across the whole spectrum of human T cell subsets. Therefore, we used in vitro models of human CD8^+^ T cell activation and exhaustion to comprehensively profile the metabolic phenotypes associated with distinct differentiation stages under controlled conditions. Combined analysis of transcriptomics and ^13^C metabolomics data revealed metabolic checkpoints in glycolysis and the TCA cycle of quiescent naive cells. Metabolic reprogramming towards aerobic glycolysis and anabolism coincided with engagement of the proline cycle for mitochondrial redox shuttling. Further differentiation towards effector cells was correlated with a progressive reliance on glycolysis. Moreover, glycolysis, mitochondrial metabolism and amino acid uptake were restricted in exhaustion. Overall, our findings extend the roles of glycolysis and mitochondrial metabolism in quiescence, activation, effector differentiation and exhaustion. Understanding how metabolic profiles differ between T cell subsets can lead to a new potential strategy in immunotherapy where T cells can be forced to adopt a stem-cell memory phenotype in order to improve anti-tumor immune responses (Dabi et al. 2022).

## Results

### In vitro activated naive CD8^+^ T cells differentiate into distinct transient subsets

To study the differentiation dynamics of human T cells, we employed an in vitro model of naïve T cell (TN) activation (Fig. 1A). Sorted CD8^+^ TN cells from three healthy donors were stimulated with anti-CD3/CD28 beads and IL-2 to drive activation and differentiation towards stem-like (TSCM) and central memory (TCM) cells, followed by restimulation to promote full differentiation of effector (TEM) cells. Based on the expression of a panel of markers, we confirmed the presence of these distinct phenotypes at different time points by flow cytometry (Fig. S1), showing that our data are consistent with a hierarchical differentiation model where the majority of activated cells acquire an effector phenotype after passing through a stem-like progenitor phase, followed by a contraction phase at the end.

**Figure 1:**
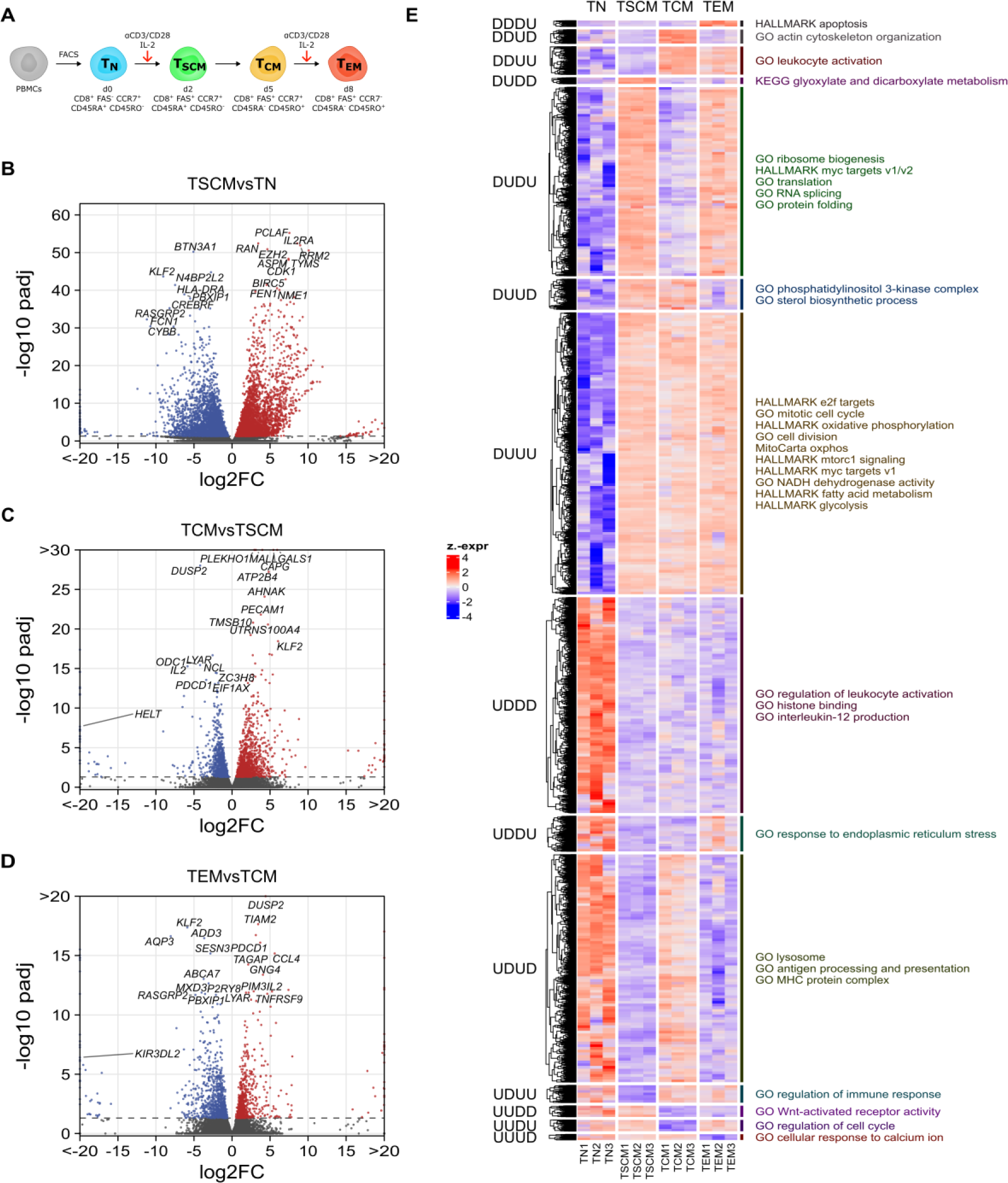
Transcriptomic characterization of CD8^+^ T cell subsets. **A.** Experimental Setup. Sorted naïve CD8^+^ T cells (TN) are stimulated with anti-CD3/anti-CD28 beads in presence of IL-2. At day 2, a population of FAS^+^ stem cell memory cells (TSCM) is established that further differentiates into a population of central memory (TCM) cells that peaks at or after day 5, coinciding with an isoform switch in CD45. Additional restimulation induces terminal differentiation and downregulation of the lymph-node homing receptor CCR7. **B-D.** Volcano plots showing differentially expressed genes (padj < 0.05) between consecutively sampled subsets. **E.** Heatmap of differentially expressed genes partitioned into patterns of up- and downregulation during differentiation (D=down, U=up), and pathway overrepresentation analysis of genes in selected clusters.

### Transcriptomic characterizations show metabolic pathway changes

Based on our initial observation of surface marker dynamics during differentiation, we characterized gene expression changes on a global scale by RNA sequencing, followed by differential gene expression analysis. Fig. S2 shows the expression levels of selected marker genes. Quiescence- (*FOXO1*, *BTG1/2*) and stemness-associated genes (*TCF7*) were highly expressed in naïve cells. Activation markers (*ILRA*, CD27, CD28) were upregulated following activation and effector proteins (*IFNG*, *GZMB*) were most strongly upregulated in TEM cells. Among inhibitory receptors, *VSIR* (VISTA) was high in TN, *PDCD1* was transiently upregulated following the initial stimulation, and both *PDCD1* and *LAG3* were upregulated following re-stimulation in TEM. Thus, we conclude that in vitro activated TN cells transiently pass through a stem-like progenitor stage, followed by central memory and effector stages before contraction.

Differential gene expression analysis revealed global transcriptomic changes between these CD8^+^ T cell subsets (Table S1), with the strongest effect following the initial stimulation of TN cells. At a significance level of 0.05, 7076 genes were differentially expressed between TN and TSCM cells (Fig. 1B), 2164 genes changed between the TSCM and the TCM subset (Fig. 1C), and 1591 differed significantly upon restimulation and further development of TCM into TEM cells (Fig. 1D). *KLF2*, which has been previously associated with quiescence (Hart, Hogquist, and Jameson 2012), was highly expressed in TN and downregulated upon activation in TSCM cells. Besides classical activation markers, genes involved in cell cycle regulation (*CDK1*, *AURKB*) and nucleotide biosynthesis (*TYMS*, *RRM2*) were among the highest upregulated genes in TSCM, along with other key metabolic genes like *GAPDH*, *ENO1*, *FABP5*, *LDHA* or *TXNIP*. Compared to TSCMs, TCMs upregulated several genes also found in TNs (*KLF2*, *SLFN5*) and downregulated several metabolic genes (*ODC1*, *NAMPT*). TEMs had high expression of genes associated with effector cells, including *PDCD1*, *IL2*, *FASLG* and *IFNG*.

As many differentially expressed genes were shared between subsets, we partitioned these genes into groups of up (U)- and downregulation (D) in different subsets and performed gene set overrepresentation analysis on the individual clusters to systematically investigate the dynamics of pathway activation (Fig. 1E, Table S2). Gene sets overrepresented only in TN cells (UDDD) included pathways involved in histone binding and modification, transcriptional repression and in the generic activation of immune cells.

Various processes related to cell cycle control, DNA replication and cell proliferation, as well as MYC, mTORC1 and E2F signaling were highly enriched in all activated subsets (DUUU). This concurred with an upregulation of metabolic processes including OXPHOS, TCA cycle, mitochondria, glycolysis, fatty acid metabolism, nucleotide biosynthesis, NADH metabolism and succinate dehydrogenase activity, indicating major metabolic reprogramming due to increased enzyme expression. Two clusters were mainly comprised of transiently activated genes that depended on acute stimulation (i.e., genes highly expressed in TSCM and TEM cells). Therein, MYC signaling, ribosome biogenesis and metabolic pathways were even further upregulated (DUDU), whereas lysosomal pathways and antigen processing were downregulated (UDUD). Wnt signaling was a key feature of TN and TSCM cells (UUDD). Specifically enriched in TSCM (DUDD) was “KEGG glyoxylate and dicarboxylate metabolism”, and downregulated genes (UDUU) were involved in leukocyte cell-cell adhesion, apoptosis and immune activation and differentiation. Genes shared by TSCM and TCM were involved in PI3K signaling and amino acid transport. Actin cytoskeleton remodeling and leukocyte migration were characteristically activated in TCMs (DDUD), and cell cycle, Wnt signaling and histone acetylation were repressed (UUDU). Genes up in the TCM and TEM subsets (DDUU) included DNA deamination (nucleotide salvage), IFN-γ response, secretion and leukocyte degranulation. Apoptosis was most strongly upregulated in TEM cells (DDDU), together with hypoxia and lymphocyte migration, while they were devoid of Ca^2+^ signaling (UUUD). Thus, these results show that in vitro differentiation reflects the main processes of in vivo immune responses (activation, proliferation, migration, contraction) and recovered their main regulatory cascades, such Wnt and Ca^2+^ signaling, and several pathways that orchestrate the metabolic reprogramming during activation and differentiation, such as mTOR and MYC.

### ^13^C tracer analysis reveals suppression of glycolysis in TN cells

As the transcriptomic analyses also indicated prominent changes in cellular metabolism, we went on to directly investigate metabolic pathway activities by performing ^13^C tracing experiments followed by LC-MS measurements. First, we generated a global qualitative map of tracer fluxes in central carbon metabolism (Fig 2). U-^13^C-glucose was readily imported and shunted into glycolysis within 24 h of labeling in all subsets, including naive T cells, resulting in full ^13^C labeling of metabolites in upper glycolysis. In TN cells, however, no labeling was detected downstream of phosphoglycerate, indicating that no glycolytic flux went into the production of phosphoenolpyruvate (pep). This was in line with a pronounced transcriptomic downregulation of *ENO1* mRNA in TN cells, which we assumed to be the major driver of the enolase reaction, as *ENO2* was only very lowly expressed (Fig. S3). Enzymes of the rate-limiting steps of glycolysis were also differentially expressed (*HK1*, *PFK*, *PK*). Whereas *PKLR* was lowly expressed in TN cells and even further downregulated upon activation, *PKM* was abundant and increased further after activation, in line with its known role to promote anabolism by increasing the availability of glycolytic intermediates (Cao, Rathmell, and Macintyre 2014). Thus, TN cells had a pronounced suppression of glycolytic flux and did not oxidize glucose in the TCA cycle, indicating that these cells used carbon sources other than glucose for OXPHOS. In contrast, activated cells showed a consistent pattern of m+2 incorporation into the TCA cycle, indicating that glycolytic flux was directed there primarily via acetyl-CoA. To a lesser extent, m+3 labeling was observable, which likely originates from anaplerotic flux via pyruvate carboxylase, and m+4 isotopomers originating from the condensation of m+2-labeled oxaloacetate with m+2-labeled acetyl-CoA in higher-order turns of the cycle.

**Figure 2:**
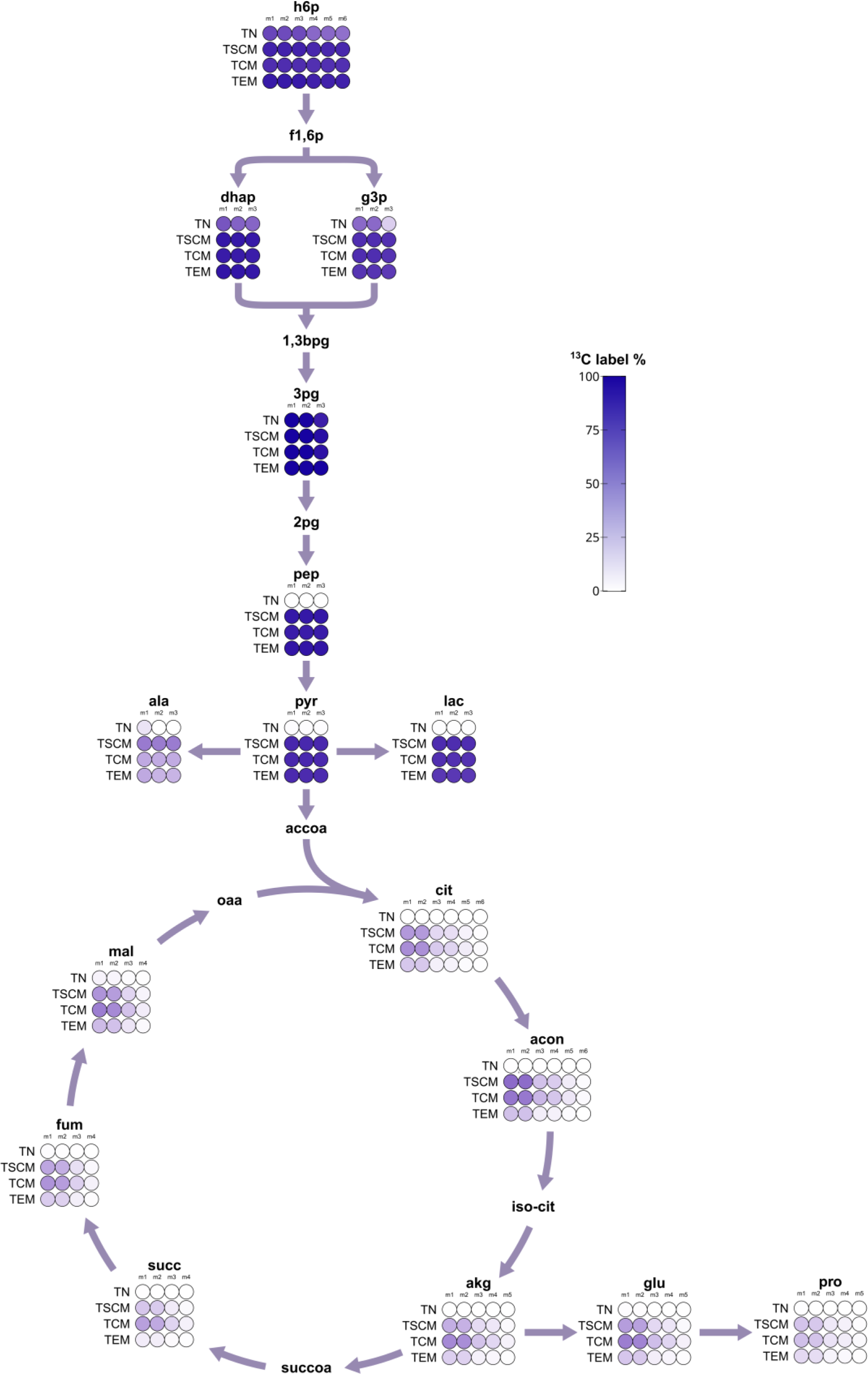
^13^C flux map of central carbon metabolism. Flux map showing the ^13^C labeling as the cumulative isotopologue distribution of selected metabolites. Color intensity indicates the fraction of labeled carbon atoms per metabolite.

### ^13^C metabolomics identifies proline cycle activity in TSCM cells

To study the metabolic dynamics of CD8^+^ T cells during activation in more detail, we performed statistical analyses of the LC-MS data to detect significant differences in both metabolite abundances (Fig 3A-C and Table S3) and labeling states (Fig 3D-F and Table S3). Compared to TN, TSCM cells had decreased intracellular levels of hexoses (hex), but increased levels of hexose-6-phosphates (h6p), glycolytic intermediates and lactate (Fig 3A). Analysis of mass isotopomer distributions further revealed significantly increased labeling of glycolytic products, in particular, a shift from unlabeled to fully labeled pyruvate and lactate, and to 50% labeled alanine (Fig 3D). As described above, the expression of glycolytic genes was increased. Thus, the suppression of glycolytic flux in naive T cells was released upon activation. Metabolites in the beginning of the TCA cycle had 30% increased labeling derived from acetyl-CoA (citrate-m+2, aconitate-m+2), which declined to 10-20% further downstream. Surprisingly among the TCA cycle metabolites, succinate levels were increased and the fumarate/succinate ratio was decreased in TNs (Fig. S4A). Expression of succinate, fumarate and malate dehydrogenases was also decreased (Fig. S5), implying substrate accumulation due to downregulation of succinate dehydrogenase (SDH) activity in TN cells. As expected, TSCM cells also upregulated several biosynthetic pathways. Aspartate, glutamate and proline exhibited the same labeling patterns as their respective precursors in the TCA cycle due to ongoing biosynthetic flux. Nucleotide, glycosylation (UDP-GlcNAc) and glucuronidation (UDP-glucuronate) substrate levels and labeling were increased and the glycine/serine ratio was higher (Fig. S4B and C), indicating increased serine biosynthesis and demand for 1C units.

**Figure 3:**
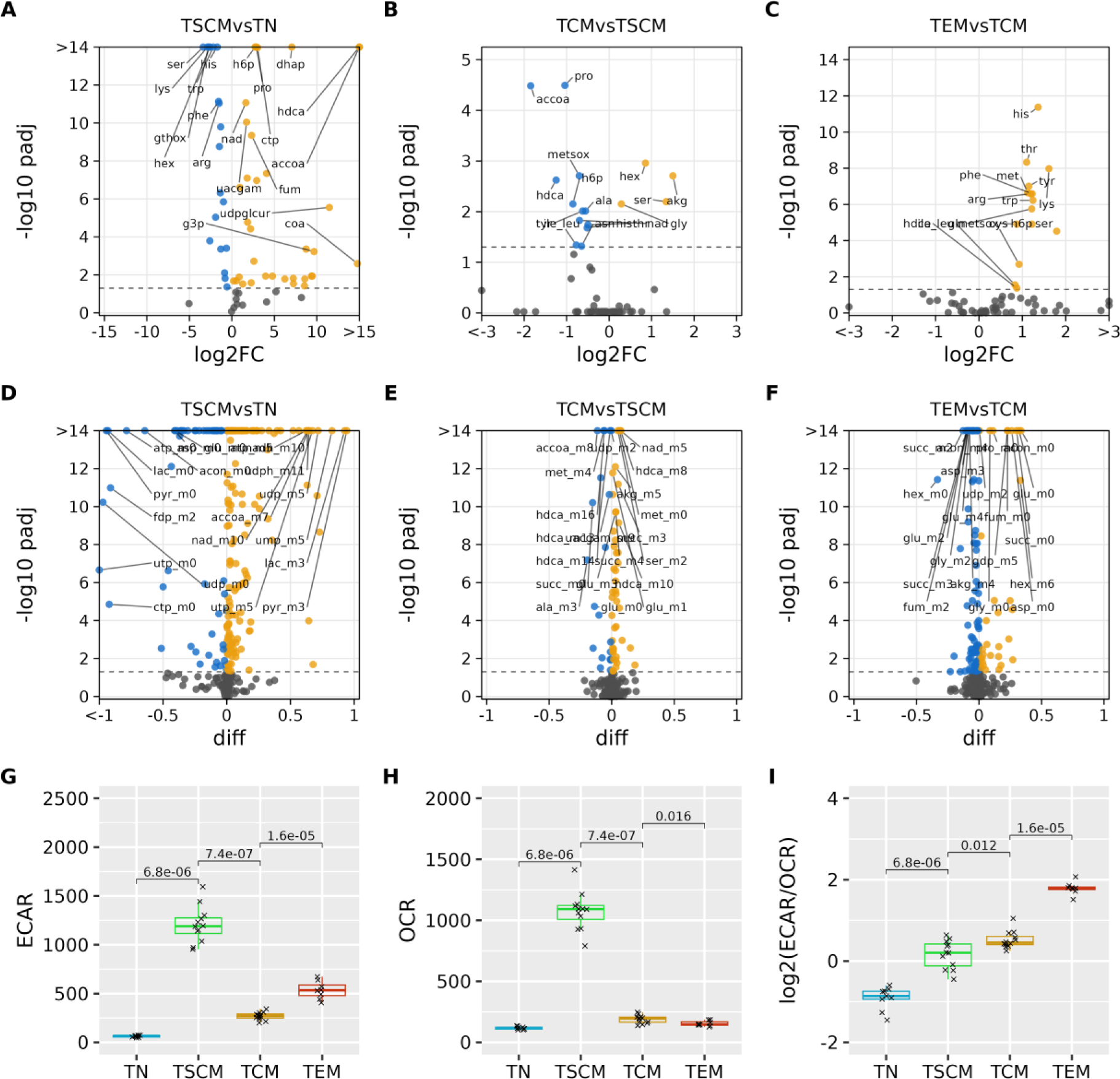
Metabolic characterization of CD8^+^ T cell subsets. Volcano plots of metabolite abundances (**A-C**) and isotopomer fractions (**D-F**). **G.** Seahorse analysis of ECAR (extracellular acidification rate, mpH min-1 10^6 cells-1) and OCR (oxygen consumption rate, pmol O2 min-1 10^6 cells-1) measured on a Seahorse analyzer, tested for differences using the Wilcoxon test.

The GSSG/GSH ratio was high in TN cells compared to all other subsets (Fig. S4D). Moreover, in TSCM cells, most amino acid levels were decreased, whereas biosynthesis and the expression of amino acid transporters (*SLC7A5*, *SLC3A2*, *SLC1A5*, *SLC38A5*) were simultaneously upregulated. Notable exceptions among amino acids were significantly higher concentrations of proline and alanine in TSCM compared to TN cells. TSCM cells also had the highest NADH/NAD+ ratio and increased labeling of NADH. Continuous regeneration of NAD+ from NADH is required to support high rates of glycolytic flux, which can be achieved in the LDH reaction together with conversion of pyruvate to lactate. As expected, we found increased *LDHA* expression (log2FC=2.7). However, we also found production of alanine from pyruvate, as well as increased flux into the TCA cycle, and both of these pathways can not be directly used to regain NAD+. Rather, catabolisation of pyruvate and subsequent oxidation of TCA cycle intermediates in mitochondria produces additional NADH from NAD+. NADH is eventually oxidized to NAD+ in OXPHOS, but then needs to be transferred across the mitochondrial membranes to be used in glycolysis. Several shuttle systems exist for this purpose. We found increased expression of genes involved in the malate-aspartate shuttle and partial upregulations of the glycerol-3-phosphate shuttle and the citrate-malate shuttle (Fig. S6). In addition, genes involved in the proline cycle were upregulated (*PYCR1*, *PYCR2* and *ALDH18A1*). As we also found an increase in proline levels and labeling, we therefore suggest that TSCM cells utilize the proline cycle to maintain NADH balance. In TCM cells, proline levels were moderately reduced compared to TSCMs (Fig. 3b), *PYCR1* was downregulated and the glycolytic intermediate dihydroxyacetone phosphate (dhap) was decreased. Labeling of TCA cycle metabolites was higher in TCM cells, most likely due to a decrease in both, draining cataplerotic reactions and replenishing anaplerosis (Fig. 3E). Decreased glutamine catabolisation is in line with a lower activity of MYC, a known driver of glutaminolysis. Compared to TCM cells, the intracellular levels of most amino acids were increased in the TEM subset (Fig. 3C), reflecting decreased consumption, as the expression of importers was only marginally increased or even decreased (Fig. S7). Whereas glycolytic intermediates were increased and the labeling remained high in glycolysis, the labeled fractions of TCA cycle intermediates were decreased (Fig. 3F). Thus, high glycolytic flux, decreased biosyntheses and decreased entry of glucose carbon into the TCA cycle were characteristic of TEM metabolism.

### Extracellular flux analysis reveals a shift to glycolysis during effector differentiation

We performed extracellular flux analysis on a Seahorse analyzer to relate the observed intracellular metabolic fluxes to extracellular acidification rates (ECAR) and oxygen consumption rates (OCR) on a global scale (Fig. 3G-H). Unstimulated TN cells had only basal ECAR and OCR. TSCMs were the most active subset and operated at maximum capacities of both ECAR and OCR. Both TCM and TEM cells had low OCRs that were only slightly higher than that of TNs, but had higher spare respiratory capacity. ECAR was similarly decreased in TCMs, but upon restimulation and effector differentiation, ECAR significantly increased to rates approximately half of those of TSCMs in the TEM cells. Despite these pronounced differences in metabolic rates between subsets, we found a progressive and gradual increase in the ECAR/OCR ratios in consecutive subsets (Fig. 3I), suggesting that CD8^+^ T cells increasingly rely on glycolysis during differentiation. To further investigate these findings on a transcriptional level, we checked the expression of OXPHOS subunits (Fig. S8). Unexpectedly, most of the respiratory complexes were not decreased or even increased in TEM. Additionally, a regression model trained on public datasets with mitochondrial copy numbers available predicted a high abundance of mitochondria in TEM cells (Fig. S9A), indicating that the observed shift towards glycolysis was not due to decreased mitochondrial abundance or changes in gene expression.

### Exhausted T cells are metabolically impaired and share features of quiescent subsets

Finally, we checked whether our observations were related not only to the differentiation but also to the proper function and dysfunction of T cells. We adopted an in vitro exhaustion model (Trefny et al. 2023) to generate control and hypofunctional T cells (Fig. 4A). Briefly, CD8^+^ T cells from healthy human donors were transduced with a T cell receptor specific for the NY-ESO-1 cancer antigen, and the transgenic T cells were stimulated with NY-ESO-1 either acutely or repeatedly to induce full effector function (TEFF) or a hypofunctional state of exhaustion (TEX), respectively. RNA sequencing analysis found 3625 differentially expressed genes between TEX and TEFF cells (Fig 4B, Table S1). Whereas most effector molecules (*GZMB*, *LTA*) were decreased, expression of *GNLY* was increased in exhausted cells. As in quiescent naive T cells, the transcription factors *KLF2*, *TCF7* and *BTG1* were upregulated. Notably, among the genes with metabolic functions, a regulator of redox metabolism and glycolysis, *TXNIP,* had the most differentially upregulated transcript in TEX. We further confirmed differential regulation of these genes (Table S4) in a previously published dataset (Trefny et al. 2023).

**Figure 4:**
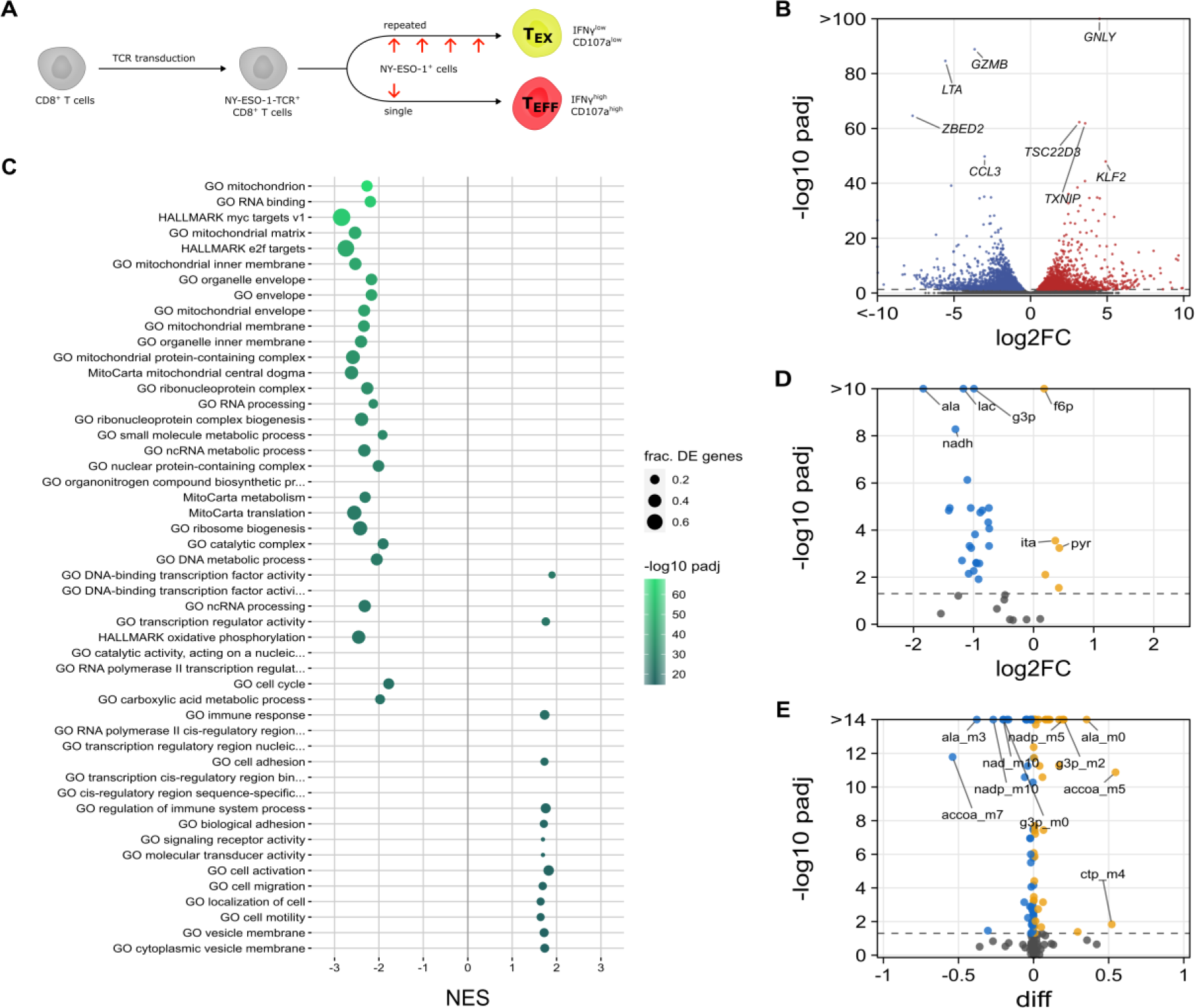
Characterization of exhausted CD8^+^ T cells. **A.** Experimental Setup. **B.** Volcano plot of differentially expressed genes. **C.** Plot of selected significantly enriched gene sets (ES > 0 up in TEX vs. TEFF, ES < 0 down in TEX vs. TEFF). Volcano plots of differentially abundant metabolites (**D**) and isotopologues (**E**).

Gene set enrichment analysis (Fig. 4C, Table S5) found an upregulation of DNA-binding transcription factor activity, with *KLF2* and *TCF7* as the top contributing genes. Also upregulated were cell adhesion terms and “GO negative regulation of leukocyte mediated cytotoxicity”. A downregulation was observed of gene sets involved in the main signaling pathways involved in T cell activation, including MYC, E2F, mTORC1 signatures. Moreover, terms related to cell cycle, ribosomes, OXPHOS, glycolysis and amino acid metabolism were downregulated. Most striking, however, was the depletion of mitochondrial components. Consistently, the predicted mitochondrial abundance was significantly decreased in TEX cells (Fig. S9B).

We further performed ^13^C tracer studies with exhausted and control effector cells (Table S6). Whereas pyruvate levels were slightly higher in TEX compared to TEFF cells, lactate levels were decreased and less labeled (Fig. 4D-E). Consistent with lower glycolytic flux, *LDHA* and key glycolytic enzymes were expressed at lower levels in TEX. Most amino acids had decreased intracellular concentrations, particularly alanine and proline. Genes of the proline cycle were strongly downregulated (*PYCR1*, *PYCR2*, *PYCR3* and *ALDH18A1*) while the NAD+/NADH ratio remained high, suggesting that TEX cells could easily maintain redox balance even under low lactate production. Alanine production is usually coupled to glycolytic flux and its labeling was likewise decreased. Presumably, TEX cells need less nutrients for protein synthesis and growth, so lower amino acid levels would hint towards decreased uptake. Indeed, we found that several of the main amino acid transporters were downregulated in TEX, including *SLC7A5*, *SLC1A5* and *SLC38A5* (Fig. S7). In the TCA cycle of TEX cells, succinate was slightly but significantly increased and the fumarate/succinate ratio was decreased (Fig. S4F), indicating lower SDH activity. Expression of SDH was moderately decreased, as well as MDH2 and, most pronouncedly, FH. Interestingly, TCA-cycle-derived itaconate levels showed a small but significant increase in TEX. Together, these data indicate that exhausted cells had reduced effector functions and partially activated signaling pathways associated with quiescent cells. TEX cells were less metabolically active than TEFF cells, had lower glycolytic flux and were depleted in intracellular amino acids due to restricted uptake. In addition, changes in the TCA cycle were associated with lower SDH activity and their mitochondrial abundance was decreased.

## Discussion

Metabolic reprogramming of CD8^+^ T cells crucially determines the outcome of immune responses to infection and cancer and has thus been investigated in great detail in mouse and human studies. However, a comprehensive characterization of metabolic fluxes in the complete course of differentiation of naive human T cells to terminally differentiated effector cells is missing. Therefore, we employed the widely used stimulation of naive T cells with anti-CD3/CD28 beads and IL-2 to differentiate them in vitro and characterize the metabolic dynamics of subpopulations following activation. Flow cytometry confirmed the presence of TSCM, TCM and TEM cells based on a panel of surface markers. Stem-like CD8^+^ T cells have been found in various in vivo studies; however, activated, highly proliferative stem-like cells in the early phases of an immune response may differ in some aspects from preferentially resting stem-like cells during memory phases after antigen clearance. Indeed, stem-like CD8^+^ T cells have been described as both proliferative (Gattinoni et al. 2009) or quiescent (Bresser et al. 2022). In acute LCMV infection, stem-like cells from early (day 8 p.i.) and late (day 30 p.i.) phases of the immune response were transcriptionally and functionally highly similar (Pais Ferreira et al. 2020), although human TSCM and TCM cells have been reported to transiently express activation and effector markers like KI67, KLRG1, PD1 or GZMB following YFV vaccination (Fuertes Marraco et al. 2022). Thus, we conclude that a transient population of activated, highly proliferative TSCM cells arises early after in vitro priming of TN cells. Without further antigenic signals, these cells further settle into a state of TCM cells that, upon restimulation, differentiate into short-lived TEM cells with full effector functions. We further found that exhausted cells displayed features of both stem-like and quiescent T cells, which is in agreement with the frequently observed negative correlation between stemness and effector functions and could adapt these cells to a continuous fight against persistent antigen without overarching inflammation.

Unstimulated naive CD8^+^ T cells displayed only basal metabolic rates and had flux constraints in lower glycolysis and the TCA cycle. This breakpoint in glycolysis was centered around enolase (ENO1) (Fig. 2), which has a critical role in supplying phosphoenolpyruvate for T cell activation. Phosphoenolpyruvate inhibits SERCA activity by promoting cysteine oxidation, thereby prolonging TCR-induced Ca2+ signaling (Ho et al. 2015). Decreased phosphoenolpyruvate levels due to glucose deprivation have thus been shown to induce T cell anergy and TIL dysfunction. Moreover, ENO1 activity was post-translationally impaired in mouse and human TILs, and this suppression was prevented by combinatorial checkpoint therapy (Gemta et al. 2019). Thus, our findings suggest that ENO1 is selectively downregulated in naive T cells to ensure suppression of activation, thereby acting as a novel quiescence checkpoint. Upon activation, cell cycle entry and differentiation, TSCM cells proliferate rapidly and turn up their biosynthetic machinery to operate at its maximum, consistent with our ECAR and OCR measurements (Fig. 3). High glycolytic flux provides intermediates for biosynthetic pathways, while a considerable fraction thereof is used to produce lactate. In the extracellular environment, exported lactate may contribute to the acidification of lymph nodes (H. Wu et al. 2020), support VISTA signaling (Johnston et al. 2019) and promote T cell stemness (Cheng et al. 2023). Intracellularly, increased lactate levels may promote lactylation (D. Zhang et al. 2019), a novel post-translational histone modification that preferentially affects metabolic enzyme expression (Yang et al. 2023). Moreover, extracellular lactate treatment has also been found to promote the stemness of CD8^+^ T cells by inhibiting histone deacetylation (Feng et al. 2022). Inhibition of LDH prevented effector differentiation and promoted stemness of CD8^+^ T cells, whereas transient inhibition allowed for full differentiation with an increased pool of stem-like cells (Hermans et al. 2020). A switch-like transition from OXPHOS to glycolysis has been consistently found in many studies of T cell metabolism (Reina-Campos, Scharping, and Goldrath 2021). Our ^13^C tracing and Seahorse data (Fig. 3) extend these findings by revealing a progressive, gradual shift towards glycolysis during differentiation, which was independent of the baseline metabolic activities and proliferation rates of the cells. Rather, the fraction of glycolytic flux that was used in OXPHOS steadily decreased. Notably, the GAPDH protein can function as a sensor of glycolytic flux that, when not bound to substrate, suppresses translation of IFN-γ mRNA in CD8^+^ T cells (Chang et al. 2013), explaining the necessity of glycolytic flux in effector cells. Interestingly, transcriptomic signatures of mitochondria were not reduced in TEM cells, suggesting that the glycolytic shift was not due to decreased mitochondrial abundance (Fig. S9A). Asymmetric inheritance of mitochondria during T cell differentiation, however, can lead to an enrichment of dysfunctional mitochondria with decreased capabilities for OXPHOS in effector cells (Adams et al. 2016). Recent investigations further suggested that mitochondrial translation, rather than metabolism, is needed for the synthesis of selected effector proteins (Lisci et al. 2021) and is enhanced during inflammation by fever (O’Sullivan et al. 2021). In contrast, exhausted cells exhibited a profound decrease in mitochondrial abundance (Fig. S9B), suggesting that a T-cell-intrinsic exhaustion program that is independent of microenvironmental stresses is sufficient to induce mitochondrial insufficiency. Exhausted T cells also had decreased glycolytic flux (Fig. 4), as previously described (Rahman et al. 2021). In addition, we found that TXNIP was increased in exhausted and also in naive cells (Elgort et al. 2010) (Levring et al. 2019). TXNIP is an endogenous inhibitor of thioredoxin-1 (TRX1) that is repressed by MYC upon T cell activation. TRX1 is critical for T cell function (Muri and Kopf 2021) and downregulated in quiescent T cells (Muri et al. 2018). To our knowledge, TXNIP has not been directly linked to T cell exhaustion. Notably, TXNIP also inhibits glycolysis (N. Wu et al. 2013), which implies an additional mechanism contributing to exhaustion that is also found in quiescent cells.

Limiting factors for biosynthesis and growth include protein synthesis, ATP production and NADH balance (Lunt and Vander Heiden 2011), all of which are upregulated in TSCM cells. Besides lactate secretion, we also found export of alanine and pyruvate, as previously found in cancer cells (DeBerardinis et al. 2007) and T cells (Xu et al. 2023), respectively. In addition to high metabolic demands for NAD+, this may restrict the ability to regenerate NAD+ via lactate dehydrogenase. We further found that TSCM cells had a low NAD+/NADH ratio (Fig. S4) and upregulated proline biosynthesis (Fig 3, Fig. S6), possibly to engage the proline cycle to transfer reducing equivalents from the cytosol to mitochondrial OXPHOS and thereby regenerate cytosolic NAD+. Notably, overexpression of *PRODH2*, a gene of the proline cycle, has been found to enhance the metabolic fitness and anti-tumor efficacy of CAR-T cells in a mouse model (Ye et al. 2022). In addition, maintenance of a sufficiently high NAD+/NADH ratio by the malate-aspartate shuttle is needed for serine biosynthesis, 1C-metabolism and nucleotide biosynthesis in CD8^+^ T cells (Xu et al. 2023), processes we found to be particularly active in TSCM cells. Thus, our results indicate that NADH balance is a limiting factor for TSCM cell metabolism and several metabolic systems, including the proline cycle, are activated to increase the capacity for NAD+ regeneration.

We further linked reduced TCA cycle flux to both quiescence and exhaustion. In particular, enzymes of later stages of the cycle were downregulated in TN cells, resulting in reduced flux through the SDH, FH and MDH reactions. SDH is necessary for T cell activation (Nastasi et al. 2021) and its downregulation may therefore represent another mechanism to ensure quiescence. Uptake of succinate can suppress T cell effector functions (Gudgeon et al. 2022), although a contrasting report suggested that cytotoxic T cells continuously need to produce and export succinate to promote autocrine signaling via SUCNR1 (Elia et al. 2022). The TCA-cycle-derived metabolite itaconate has been reported to inhibit SDH activity (Lampropoulou et al. 2016). Although we failed to detect significant differences in itaconate levels during differentiation, we found labeling patterns of itaconate that matched TCA cycle compounds in several T cell subsets, and an increase in itaconate abundance in TEX cells (Fig. 4). While initial investigations did not find any effects in CD8^+^ T cells (Nastasi et al. 2021), a more recent study reported suppressive effects of external itaconate, leading to succinate accumulation and reduced proliferation (Zhao et al. 2022). In addition, itaconate may also suppress glycolysis through post-translational mechanisms (Qin et al. 2019), consistent with our observation of decreased glycolysis in exhausted cells. These results therefore suggest that itaconate also has a T-cell-intrinsic role and may contribute to exhaustion by endogenous production.

Notwithstanding the profound impact of the extracellular microenvironment on T cell function and dysfunction, our results highlight the landscape of intrinsic regulatory points of CD8^+^ T cell metabolism in various differentiation states. Modulation of metabolic programs by activating or inhibiting these targets, even if only transiently, has the potential to arrest differentiation or promote subset conversion. The feasibility of this approach has previously been demonstrated for mTOR (Scholz et al. 2016) or Wnt inhibition (Gattinoni et al. 2009), nutrient restriction (Vodnala et al. 2019) or lactate dehydrogenase inhibition (Hermans et al. 2020). Thus, these results may provide onset points in various therapeutic settings, such as the ex vivo conditioning of T cells in cellular therapies for the treatment of cancer.

## Methods

### Cell culture

Leukoreduction system chambers (LRSCs) (Néron et al. 2007) from apheresis platelet donors were used as source of peripheral blood mononuclear cells (PBMCs) and were obtained from the Central Institute for Blood Transfusion and Immunology situated at the Tirol Kliniken GmbH in Innsbruck, Austria. At the time of the donation, platelet donors were in a healthy state and fulfilled the general requirements for donating blood in Austria. LRSCs from platelet donors were provided on informed consent after finishing the apheresis procedure. PBMCs were isolated by density gradient centrifugation on a Ficoll-plaque by using Lymphocyte Separation Medium (LSM-A) (Capricorn Scientific, Ebsdorfergrund, Germany). After Ficoll gradient separation, the PBMC fraction was collected and washed twice with ice-cold DPBS (Gibco/Thermo Fisher Scientific). PBMCs were counted under a hemocytometer and the viability was assessed via Trypan blue (Sigma Aldrich, St. Louis, USA). The cells were cultured in RPMI 1640 medium containing 10% Fetal Bovine Serum (FBS), 2 mM L-Glutamine and 1% Penicillin Streptomycin (all from Sigma Aldrich, St. Louis, USA) and incubated overnight at 37°C and 5% CO_2_ before starting the CD8^+^ T cell differentiation.

### Cell sorting

Naïve CD8^+^ T cells were sorted using a FACSAria I Sorter (BD Biosciences) by staining the isolated PBMCs with anti-CD8 FITC (BD Biosciences), anti-CD197 (CCR7) PE, anti-CD95 APC (both from BioLegend), anti-CD45RO PE-Cy7, anti-CD45RA BV421 and 7-Amino-Actinomycin D (7-AAD) (all from BD Biosciences). PBMCs were sorted into naïve T cells (TN) (CD8^+^ CD197^+^ CD95^-^ CD45RO^-^ CD45RA^+^), collected into RPMI 1640 medium containing 1% FBS and counted manually before starting the differentiation experiments.

### In vitro differentiation model

For in vitro differentiation, naïve CD8^+^ T cells (1.5 x 10^6^ cells/ml and 1 x 10^6^ cells/ml) were distributed into four wells of a 24-well plate and activated for 8 days by adding prewashed anti-CD3/CD28 Dynabeads at a 1:1 bead-to-cell ratio (Invitrogen/Thermo Fisher Scientific, Massachusetts, USA) and human rIL-2 (30 U/ml) (Roche/Merck, Darmstadt, Germany) to the culture. To generate effector memory T cells, 1 x 10^6^ cells/ml were restimulated on day 7 by adding prewashed anti-CD3/CD28 Dynabeads at a 1:1 bead-to-cell ratio and human rIL-2 (30 U/ml) a second time to the culture. The cells were cultured in RPMI 1640 medium without glucose (Gibco/Life Technologies, Carlsbad, USA) where 11 mM D-Glucose was added to 10% Fetal Bovine Serum (FBS), 2 mM L-Glutamine and 1% Penicillin Streptomycin (all from Sigma Aldrich, St. Louis, USA). 24 hours before collection, culture medium was replaced with RPMI 1640 medium containing 11 mM of ^13^C labeled-Glucose (Cambridge Isotope Laboratories, Massachusetts, USA). At indicated time points, naïve T cells (TN, day 1), stem cell memory T cells (TSCM, day 2), central memory T cells (TCM, day 5) and effector memory T cells (TEM, day 8) were collected and anti-CD3/CD28 beads were removed by using a DynaMag-2 magnet (Thermo Fisher Scientific, Massachusetts, USA) before further processing for flow cytometry, RNA sequencing and metabolomics analyses.

### Flow cytometry

To analyze cell surface marker during differentiation, 1 x 10^5^ PBMCs were labeled at every time point (day 1, 2, 5 and 8) with the same panel of human antibodies as used for cell sorting and analyzed by flow cytometry on a FACS Fortessa (BD Biosciences). UltraComp eBeads (Invitrogen/Thermo Fisher Scientific, Massachusetts, USA) were used for the compensation setup. The cell type fractions were determined as the following: Naïve T cells (TN) (CD8^+^ CD197^+^ CD95^-^ CD45RO^-^ CD45RA^+^), stem cell memory T cells (TSCM) (CD8^+^ CD197^+^ CD95^+^ CD45RO^-^ CD45RA^+^), central memory T cells (TCM) (CD8^+^ CD197^+^ CD95^+^ CD45RO^+^ CD45RA^-^) and effector memory cells T cells (TEM) (CD8^+^ CD197^-^ CD95^+^ CD45RO^+^ CD45RA^-^). Analysis was performed using FlowJo software.

### In vitro exhaustion model

In vitro exhausted CD8^+^ T cells were generated as previously described (Trefny et al. 2023). Briefly, CD8^+^ T cells isolated from healthy donor PBMCs were stimulated by a 1:1 ratio of anti-CD3/28 activation beads (Miltenyi) in complete human T cell medium (RPMI1640 from Sigma containing 2 mM glutamine with the addition of 1 mM pyruvate, 1% penicillin-streptomycin, 10% heat-inactivated AB+ male human serum, 50 nM beta-mercaptoethanol in presence of 150 U/ml rhIL2 (Proleukin). The day after stimulation, cells were transduced with VSVg-pseudotyped Lentivirus encoding a human T cell receptor specific for NY-ESO-1 (gift from Natalie Rufer and Michael Hebeisen) (Schmid et al. 2010). To generate exhausted T cells, TCR^+^ cells were stimulated every 3 days for four cycles with the HLA-A2^+^ T2 tumor cell line loaded with 1000 nM NY-ESO-1 SLLMWIQV peptide at a 1:3 effector to target ratio in T cell medium with 50 U/ml IL-2. For functional effector cells, the T cells were expanded in the same media for 9 days before a single stimulation with the peptide loaded T2 cells as described above. 3 days post last stimulation, the T cells were sorted for live TCR^+^ CD8^+^ CD19^-^ CD14^-^ CD56^-^ cells and resuspended in RPMI 1640 medium containing 5 mM of ^13^C labeled-Glucose at 1×10^6^ cells/ml with the addition of 50 U/ml IL-2. Cells were cultured in a 24-well plate for 24 h at 37°C with 5% CO_2_ before further processing for RNA sequencing and metabolomics analyses.

### Seahorse XFp Cell Energy Phenotype Test

Naïve T cells (TN), stem cell memory T cells (TSCM), central memory T cells (TCM) and effector memory T cells (TEM) were collected, counted, and centrifuged at 1500 rpm for 10 minutes at room temperature. Cells were resuspended at a concentration of 8×10^6^ cells/ml in Seahorse XF Base Medium (Agilent Technologies, Santa Clara, CA) supplemented with 1 mM pyruvate, 2 mM glutamine, 10 mM glucose (all from Sigma Aldrich, St. Louis, USA), and pH was adjusted to 7.4. Seahorse XFp Cell Culture Miniplates (Agilent Technologies, Santa Clara, CA) were precoated with poly-L-lysine (Sigma) and 1×10^5^ to 4×10^5^ cells were plated into each well. Miniplates were centrifuged at 300 g for 1 minute with low-brake deceleration. Seahorse XFp Cell Energy Phenotype Tests (Agilent Technologies, Santa Clara, CA) were performed according to the manufacturer’s protocol on an XFp instrument and data analysis was performed using the Seahorse XFe Wave software.

### Metabolomics sample preparation

After 24h culturing in ^13^C-containing medium, 1 x 10^6^ cells were collected at indicated time points and centrifuged at 2000 rpm for 1 to 3 min at 4°C. For medium extraction, 10µl of the supernatant was added to 990µl 80% methanol containing 2 µM of d27 myristic acid (pre-chilled to −80°C) (extraction buffer provided by the VIB Metabolomics Expertise Center). For cellular extraction, the supernatant was removed, the pellet was carefully washed with 1 ml of ice-cold 0.9% NaCl (Sigma Aldrich, St. Louis, USA) and this solution was then aspirated. Pellet was resuspended in 250 µl (TEFF and TEX) or 300 µl (TN, TSCM, TCM and TEM) of 80% methanol containing 2 µM of d27 myristic acid (pre-chilled to −80°C) and pulse vortexed three times for five seconds. Samples were stored at −80°C for 24 hours, centrifuged at 1500 rpm for 15 minutes at 4°C and 250 µl of metabolite-containing supernatants of each sample were sent for analysis.

### LC-MS metabolomics measurements

10 µl of each sample was loaded into a Dionex UltiMate 3000 LC System (Thermo Scientific Bremen, Germany) equipped with a C-18 column (Acquity UPLC -HSS T3 1. 8 µm; 2.1 x 150 mm, Waters) coupled to a Q Exactive Orbitrap mass spectrometer (Thermo Scientific) operating in negative ion mode. A step gradient was carried out using solvent A (10 mM TBA and 15 mM acetic acid) and solvent B (100% methanol). The gradient started with 5% of solvent B and 95% solvent A and remained at 5% B until 2 min post injection. A linear gradient to 37% B was carried out until 7 min and increased to 41% until 14 min. Between 14 and 26 minutes the gradient increased to 95% of B and remained at 95% B for 4 minutes. At 30 min the gradient returned to 5% B. The chromatography was stopped at 40 min. The flow was kept constant at 0.25 ml/min and the column was placed at 40°C throughout the analysis. The MS operated in full scan mode (m/z range: [70.0000-1050.0000]) using a spray voltage of 4.80 kV, capillary temperature of 300°C, sheath gas at 40.0, auxiliary gas at 10.0. The AGC target was set at 3.0E+006 using a resolution of 140000, with a maximum IT fill time of 512 ms. Data collection was performed using the Xcalibur software (Thermo Scientific). The data analyses were performed by integrating the peak areas (El-Maven - Polly - Elucidata).

### Metabolomics data analysis

Quantification thresholds were defined for each batch from three blank measurements as two standard deviations above blank means. Isotopologue intensities below blank means were set to zero and isotopologue intensities above blank means but below quantification thresholds were corrected by a linear mapping between zero and the quantification threshold. Only isotopologues with at least two biological replicates above quantification thresholds were used for statistical testing. Correction for natural isotope abundance was applied with the IsoCorrectoR R package 1.16.0 (Heinrich et al. 2018) in high-resolution mode. Corrected data were normalized between isotopologues by mean metabolite abundance and between samples by size factors consisting of total isotopologue sums weighted by their inverse relative variances. Isotopologue abundances were further cleaned from sample groups with more than 1/3 of the respective values below quantification threshold. Metabolite abundances were calculated from the sums of isotopologue abundances and fractions of isotopologues relative to isotopologue sums were used as mass isotopomer distributions. For differential abundance analysis, metabolite abundances were assessed in a linear mixed model using the nlme R package 3.1-162 and isotopologue fractions were modeled with beta regression using the glmmTMB R package 1.1.7. Cell type was treated as fixed and donor as random effect. The multcomp R package 1.4-23 was used for statistical hypothesis testing of particular contrasts and the resulting p-values were FDR-adjusted. Statistical analyses of isotopologue abundances using a linear mixed model and of fractional metabolite labeling using beta regression yielded similar conclusions and were thus not further discussed.

### RNA sequencing

Naïve T cells (TN), stem cell memory T cells (TSCM), central memory T cells (TCM), effector memory T cells (TEM), full effector function T cells (TEFF) and exhausted T cells (TEX) were collected, snap frozen and total RNA was isolated using the RNeasy Plus Mini Kit (Qiagen) or RNeasy Plus Micro Kit (Qiagen GmbH-Austria, Hilden, Germany), including DNAse treatment and following the manufacturer’s instructions. All Isolated total RNA samples were quality validated and submitted to library preparation following the Lexogen QuantSeq 3’mRNA protocol (Lexogen GmbH, Vienna, Austria). The resulting libraries were multiplexed and sequenced with Ion Proton technology and Ion Hi-Q chemistry (Ion Torrent, Thermo Fisher Scientific, Vienna, Austria). CUTADAPT 4.0 was used to trim sequencing reads of low-quality bases and poly-A tails with the options -q 20, -O 10, -e 0.15, -m 10 and the adapter sequence -a A100. We used the nf-core rnaseq pipeline 3.6 (10.5281/zenodo.6327553) with the trimming step disabled for the alignment of reads with STAR 2.7.9a to the human genome (GRCh38) and for the quantification of reads with SALMON 1.5.2. The correct pairing of samples was confirmed by grouping samples to donors of origin with NGSCheckMate 1.0.0. Raw counts were imported into R 4.1.3 and DESeq2 1.38.3 was used with IHW 1.26.0 to test for differentially expressed genes. For analyses of biological functions, we compiled a collection of gene sets from gene ontology (GO), HALLMARK, KEGG and the metabolic databases MitoCarta 3.0 (Rath et al. 2021) and MetabolicAtlas 3.3 (Robinson et al. 2020). For the differentiation samples (TN, TSCM, TCM and TEM), significantly different genes with padj < 0.05 were partitioned into groups of up- and downregulation according to their z-scaled expression values. The resulting gene clusters were further used for gene set overrepresentation analysis (ORA) with the ClusterProfiler R package 4.6.2. For samples from the exhaustion experiments, we performed gene set enrichment analysis (GSEA) with ClusterProfiler on genes ranked by the Wald test statistic from DESeq2. The regression model for mitochondrial abundance prediction was based on public data from (Yuan et al. 2020) and (Reznik et al. 2016).

## Data and code availability

Code to reproduce the analyses is available at https://github.com/icbi-lab/kirchmair_2023 and functions for the analysis of ^13^C metabolomics data are available as an R package at https://github.com/AlexanderKirchmair/c13ms.

## Acknowledgements

We would like to thank Camila T. Cologna from the VIB-KU Leuven Metabolomics Expertise Center for her help with metabolomics sample preparation and LC/MS-measurements.

## Supplementary Information

**Supplementary Figure 1:**
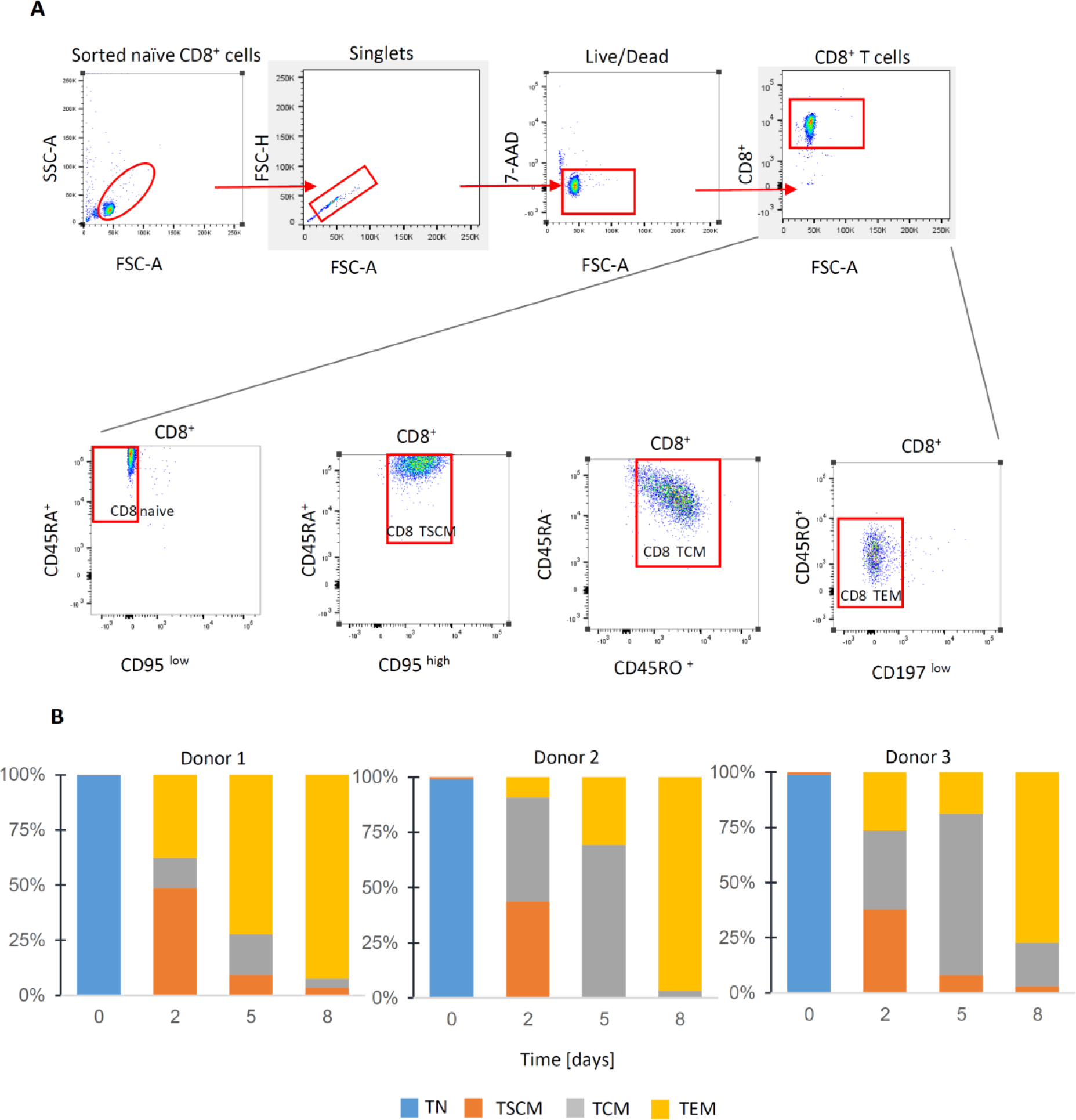
Flow cytometry. **A.** Gating strategy to identify differentiation stages of CD8+ T cells. Naïve CD8^+^ T cells were identified as CD8^+^ CD197^+^ CD95^-^ CD45RO^-^ CD45RA^+^ lymphocytes and sorted (upper left). Differentiation stages were defined based on the differential expression of CD95, CD45RA, CD45RO and CD197 on CD8^+^ T cells (lower panel). TN (naïve T cells) (CD8^+^ CD197^+^ CD95^-^ CD45RO^-^ CD45RA^+^); TSCM (stem cell memory T cells) (CD8^+^ CD197^+^ CD95^+^ CD45RO^-^ CD45RA^+^); TCM (central memory T cells) (CD8^+^ CD197^+^ CD95^+^ CD45RO^+^ CD45RA^-^); TEM (effector memory T cells) (CD8^+^ CD197^-^ CD95^+^ CD45RO^+^ CD45RA^-^). **B.** Differentiation dynamics. Changes in subset composition following 8 days of differentiation of TN cells.

**Supplementary Figure 2:**
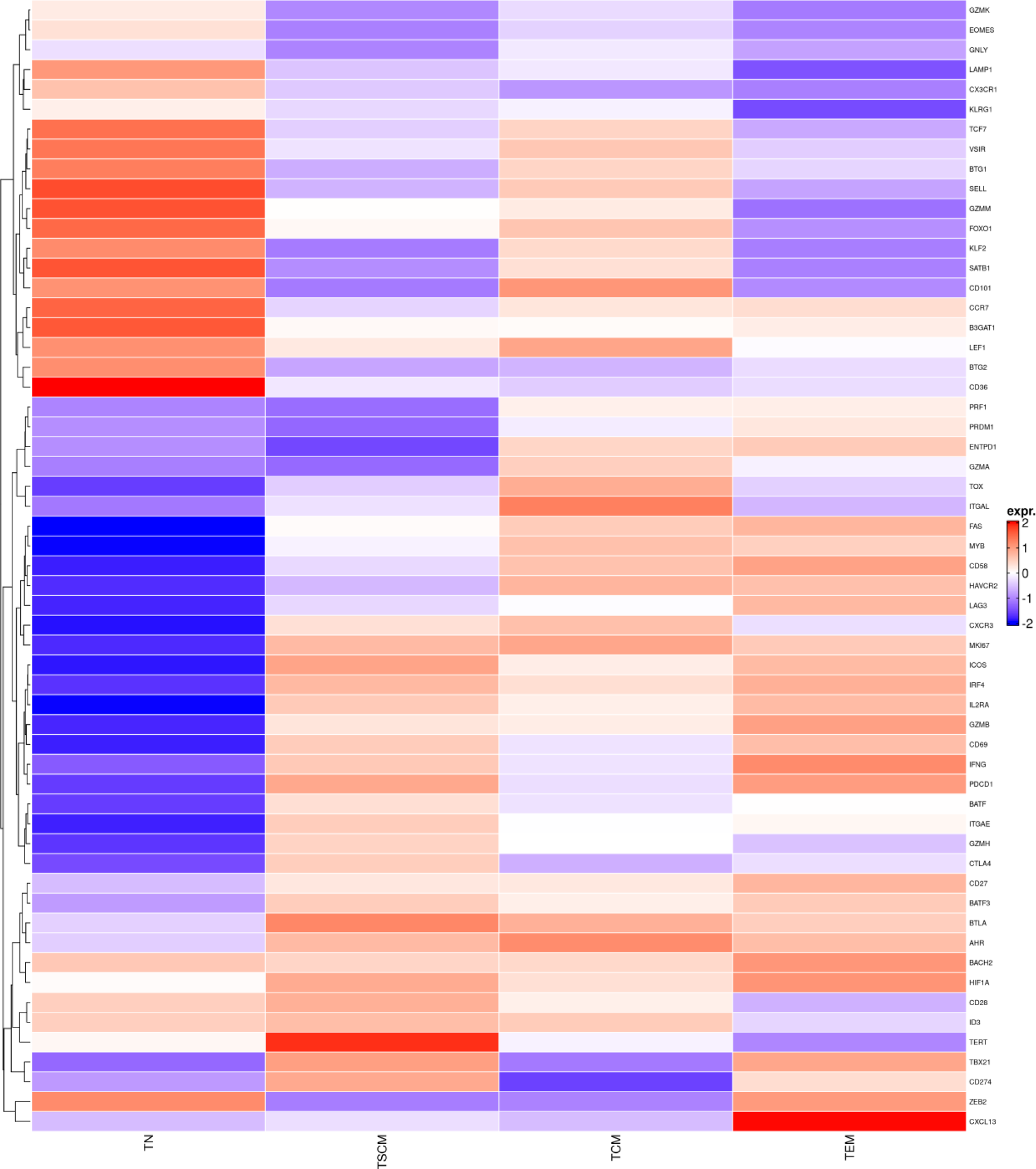
Expression of selected T cell markers.

**Supplementary Figure 3:**
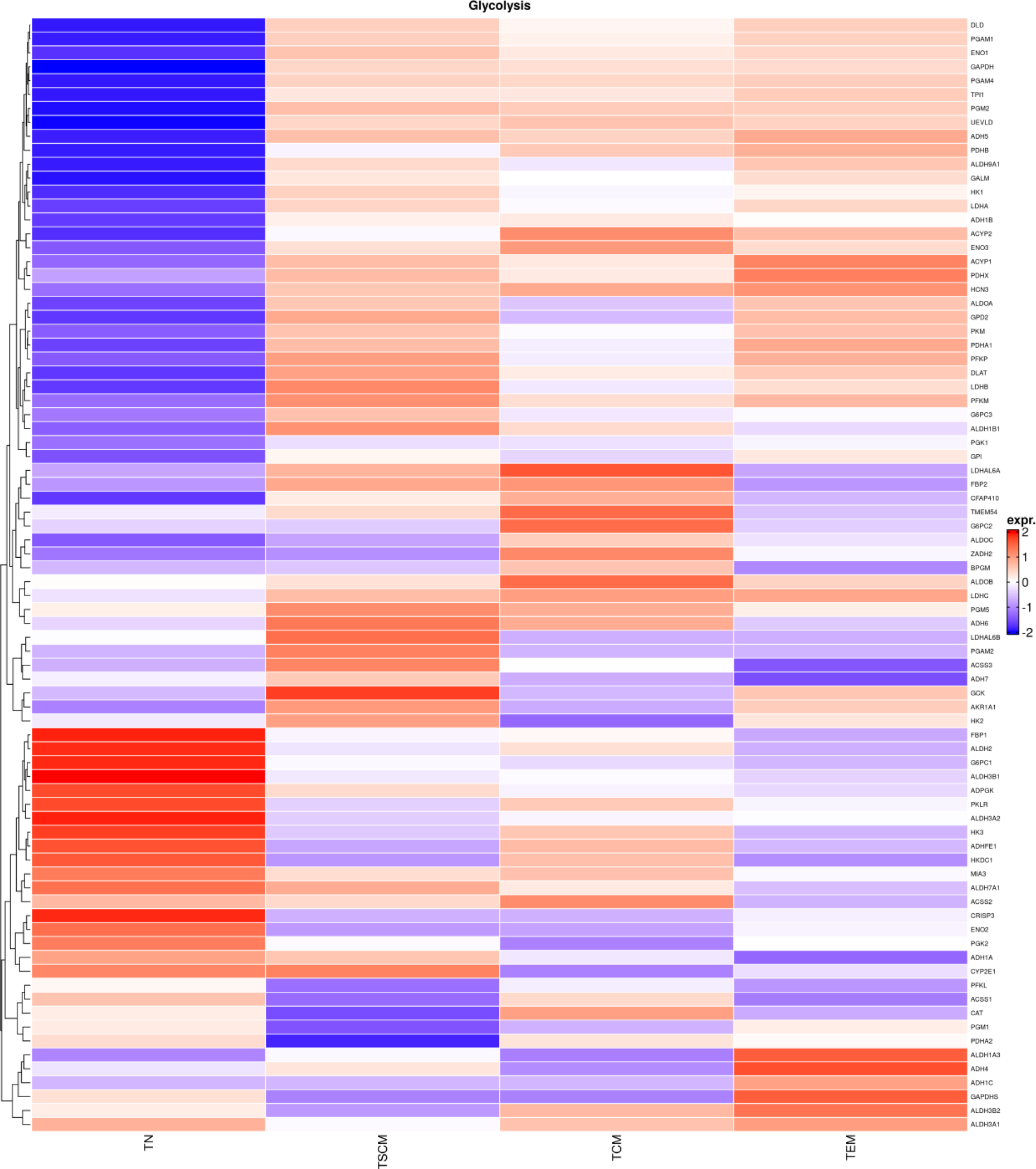
Expression of genes related to glycolysis.

**Supplementary Figure 4:**
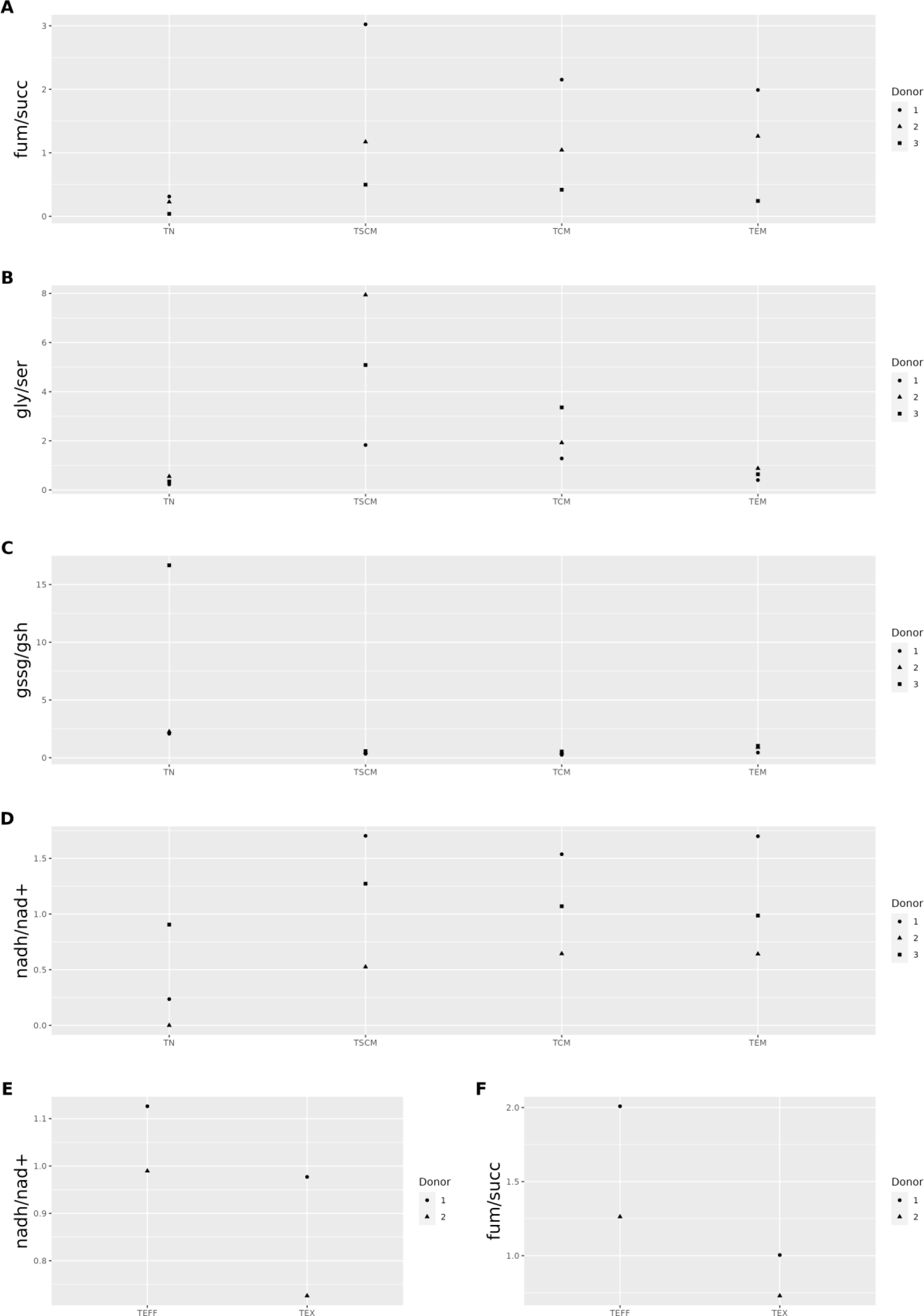
Abundance ratios of selected metabolites.

**Supplementary Figure 5:**
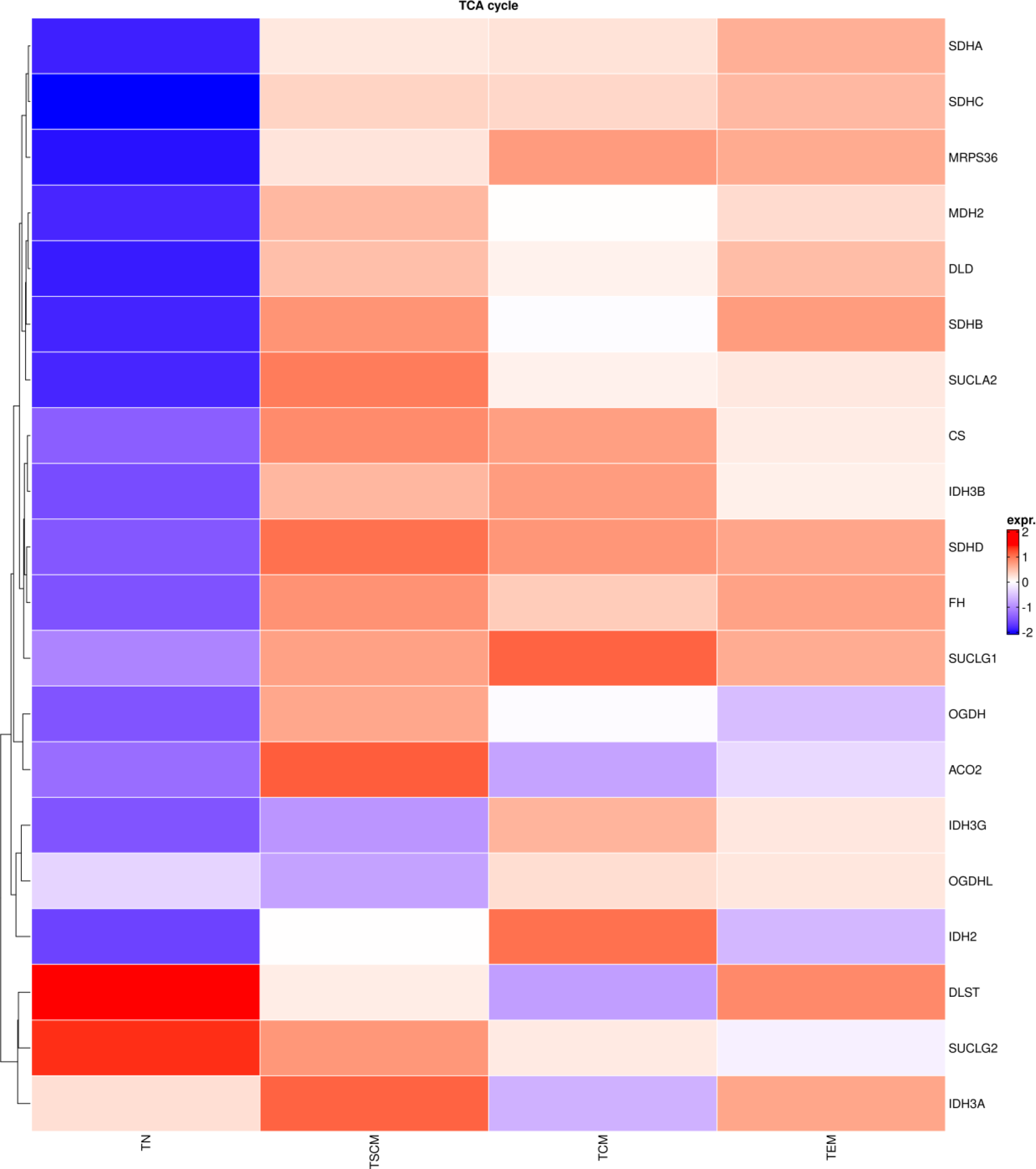
Expression of genes in the TCA cycle.

**Supplementary Figure 6:**
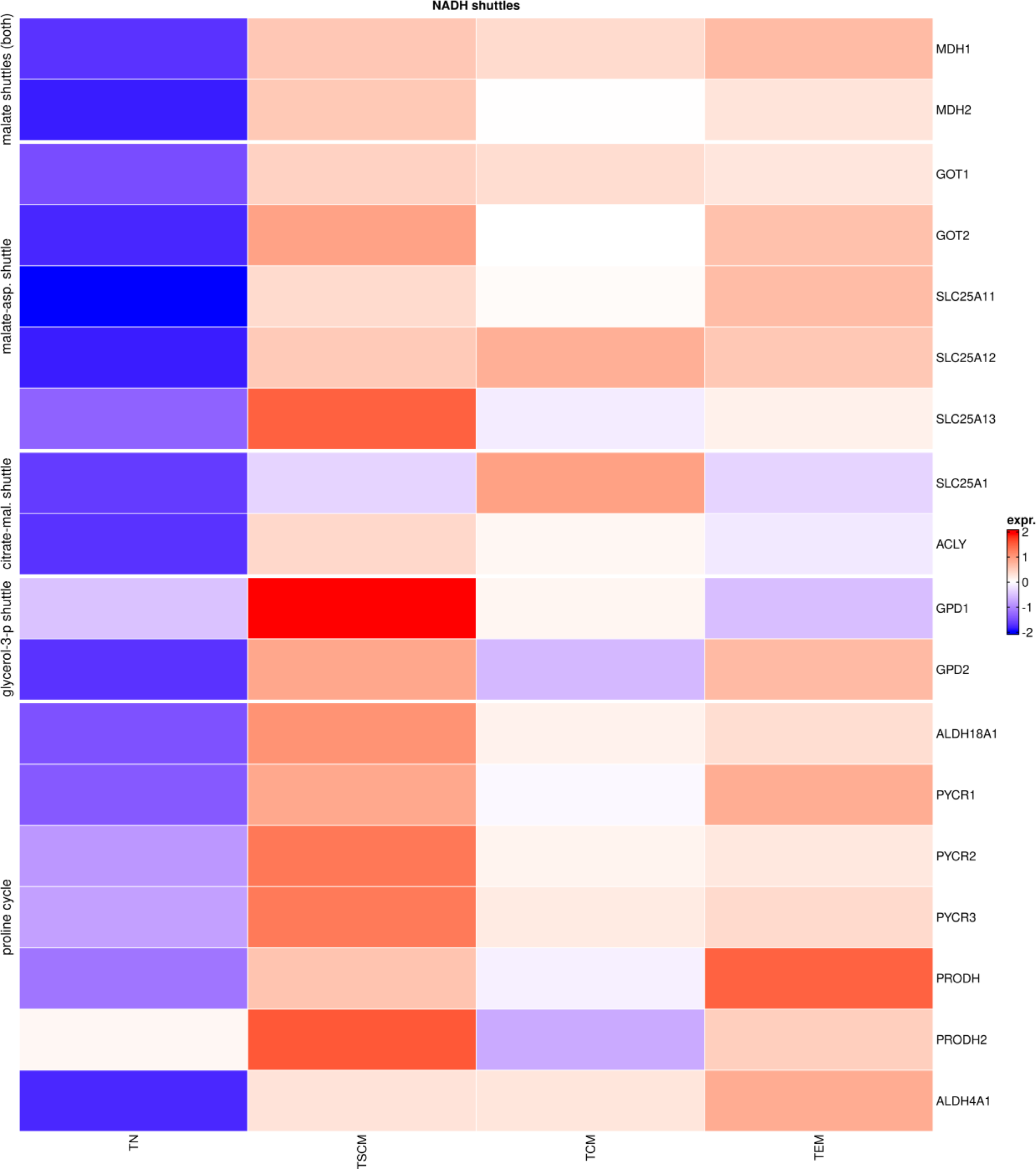
Expression of mitochondrial NADH shuttles.

**Supplementary Figure 7:**
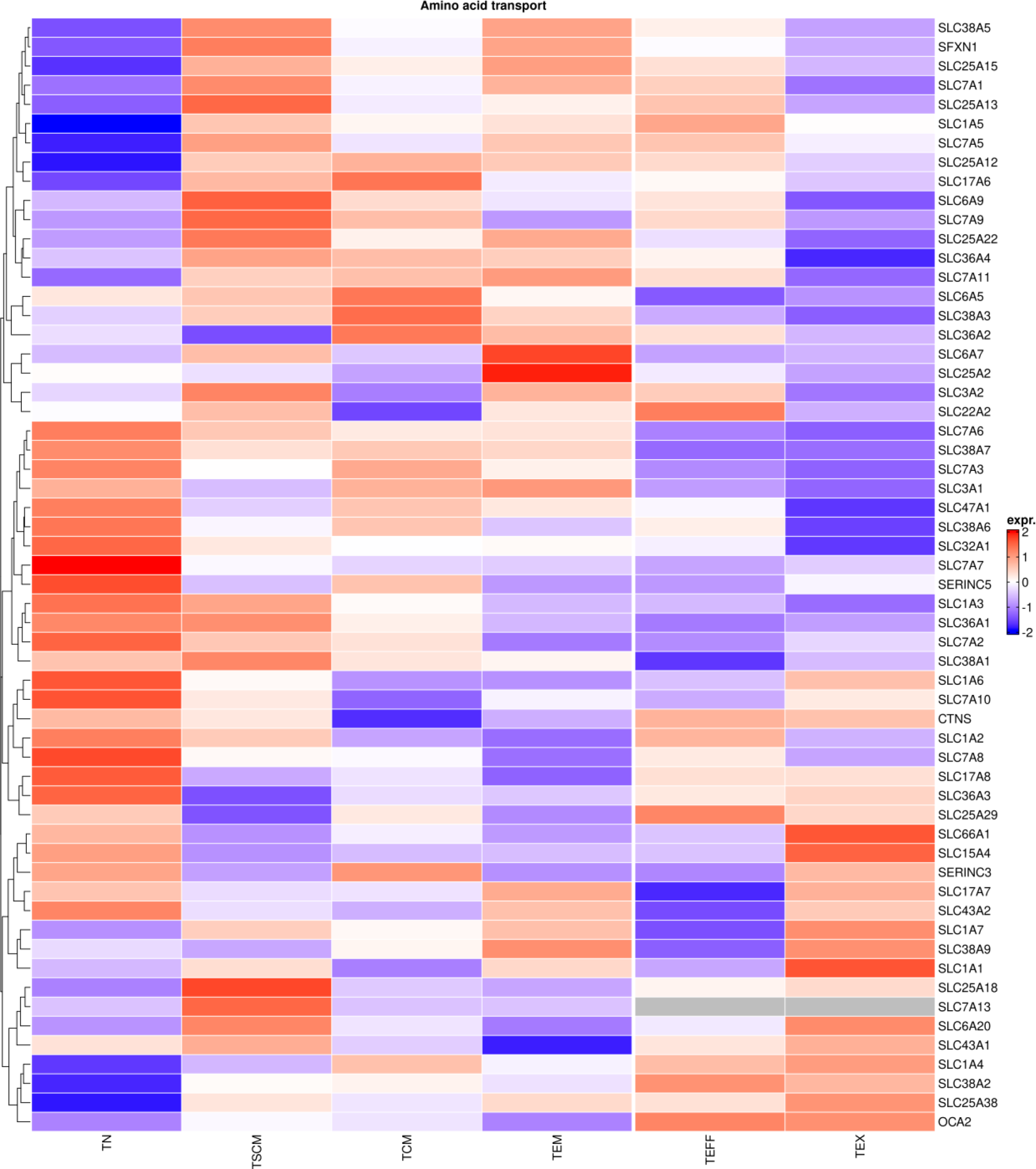
Expression of nutrient transporters.

**Supplementary Figure 8:**
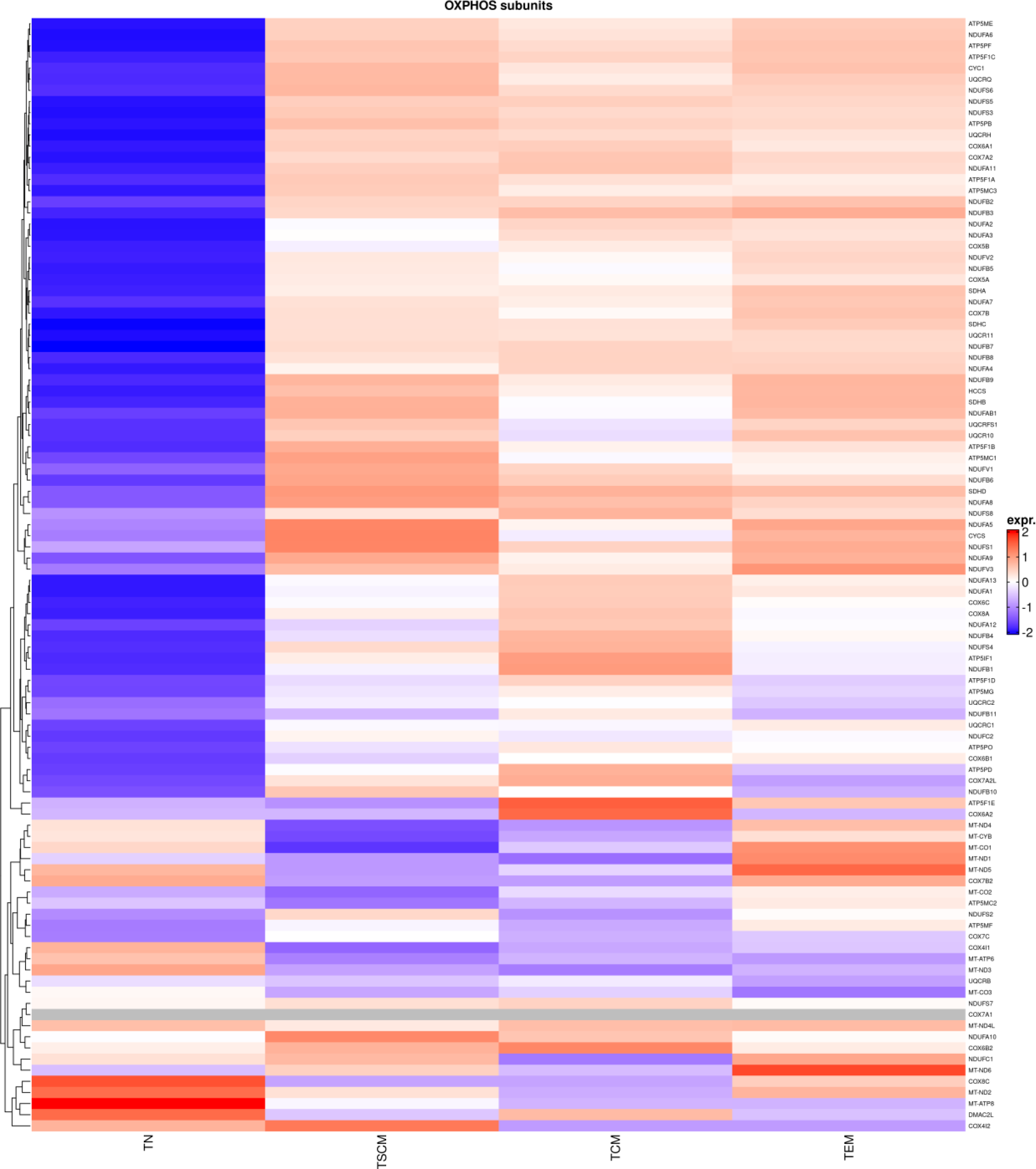
OXPHOS subunit expression.

**Supplementary Figure 9:**
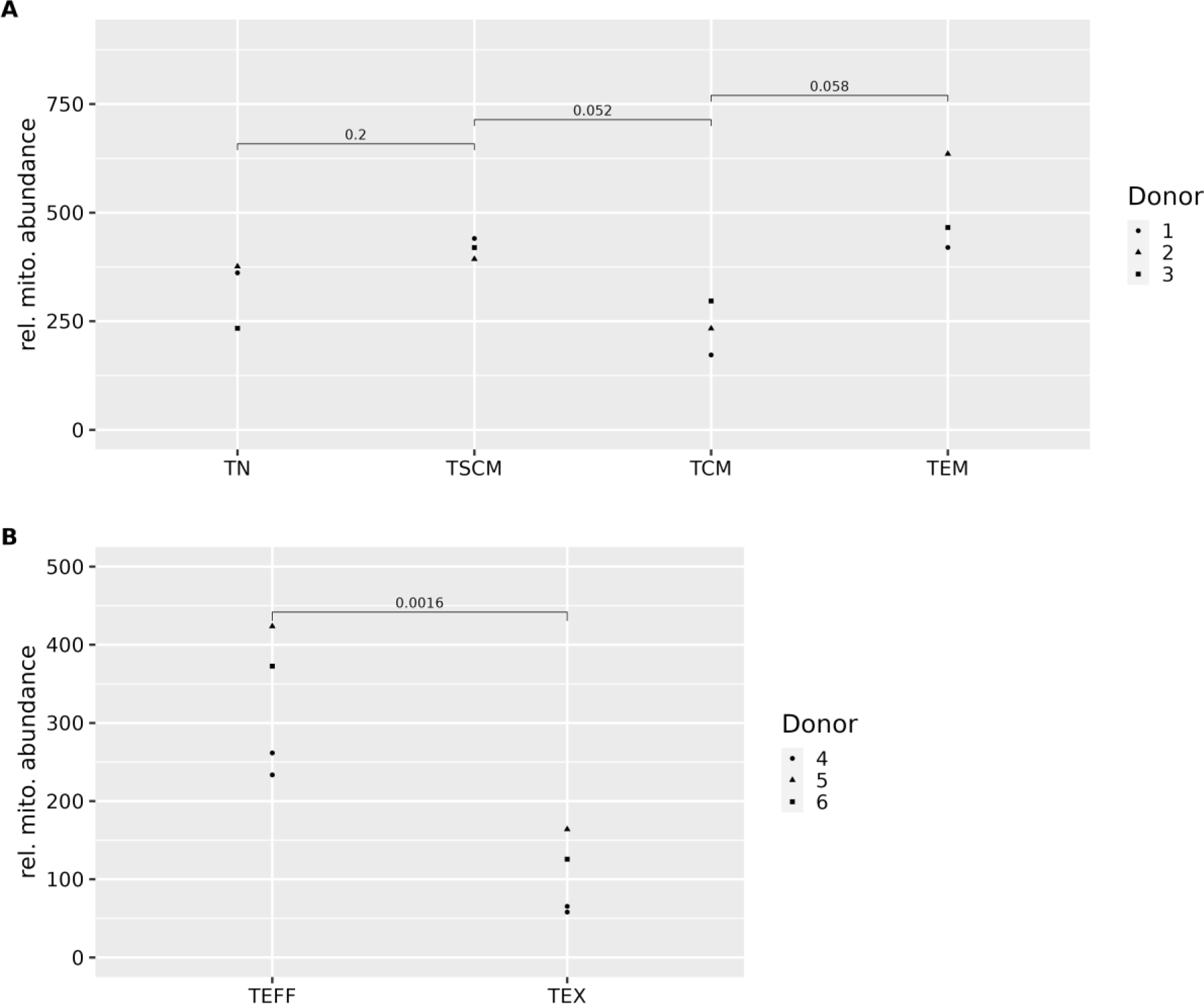
Mitochondrial gene expression. Expression of genes that correlate with mitochondrial copy numbers in public datasets, tested for differences using a paired t-test.

